# TDP-43 expression in the cytoplasm leads to early synaptic and mitochondrial abnormalities in an inducible mouse model of ALS/FTD

**DOI:** 10.1101/2025.10.01.679786

**Authors:** Florencia Vassallu, Milagros López, Florencia López Ambrosioni, Juan Casal, Laura Caltana, Lionel Muller Igaz

## Abstract

TDP-43 proteinopathy is the primary pathology associated with amyotrophic lateral sclerosis (ALS) and frontotemporal dementia (FTD), indicating that these neurodegenerative diseases have common underlying mechanisms. We have previously shown that transgenic (Tg) mice conditionally overexpressing a cytoplasmic form of human TDP-43 protein (TDP-43-ΔNLS) in forebrain neurons replicate key features of FTD/ALS, including altered cognitive, motor and social behaviors. These behavioral phenotypes and changes in plasticity-related gene expression can be detected as early as 1 month after Tg induction, before overt neurodegeneration occurs. To assess early ultrastructural features in this model, we performed Transmission Electron Microscopy (TEM) analysis in the cortex (Ctx) and hippocampus (Hp) of Tg animals and their non-Tg controls. TEM evaluation of Ctx and Hp revealed that synaptic density was significantly decreased and synapse length was increased in both regions of Tg animals. Synaptic cleft thickness was increased and post-synaptic density thickness was decreased only in the Ctx of Tg mice, revealing differential regional effects in synaptic morphology. We analysed mitochondrial density and we found an increase in the Ctx and a decrease in the Hp of Tg animals, with preserved individual mitochondrial area. Lastly, transcriptomic and proteomic analysis from both transgenic TDP-43-ΔNLS mice and human proteinopathy showed widespread decreased expression of synaptic structure and function genes. The alterations in synaptic density and architecture reported here, combined with the mRNA/protein expression data, suggest that TDP-43-ΔNLS mice may exhibit abnormal synaptic transmission and that ultrastructural changes play a role in the early behavioral deficits observed in this model.

**Highlights:**

- Cytoplasmic TDP-43 expression *in vivo* causes early synaptic and mitochondrial abnormalities.
- Reduced synaptic density observed in cortical and hippocampal regions of TDP-43-ΔNLS mice.
- Synaptic ultrastructure altered, including increased cleft width and reduced PSD thickness.
- Region-specific mitochondrial density changes: increased in cortex, decreased in hippocampus.
- Findings link TDP-43 mislocalization to early structural and functional brain deficits.

## 1. Introduction

Neurodegenerative diseases (NDs) are a heterogeneous group of disorders characterized by the progressive loss of specific neuronal populations, leading to impairments in cognition, motor function, general behavior, and ultimately, survival. The global prevalence of NDs is increasing, particularly in aging populations, posing significant challenges to healthcare systems worldwide (Prince, 2015).

A common pathological hallmark of many NDs is the accumulation of misfolded or aggregated proteins within neurons. These proteinopathies disrupt cellular homeostasis and are associated with oxidative stress, mitochondrial dysfunction, impaired proteostasis, neuroinflammation, and synaptic loss (Soto and Pritzkow, 2018). Notable examples include α-synucleinopathies such as Parkinson’s disease, tauopathies like Alzheimer’s disease (AD), and TDP-43 proteinopathies like amyotrophic lateral sclerosis (ALS) and frontotemporal dementia (FTD) (Neumann et al., 2006; Uytterhoeven et al., 2025; Vekrellis et al., 2024; Waheed et al., 2023).

TAR DNA-binding protein 43 (TDP-43) is a ubiquitously expressed RNA-binding protein involved in various aspects of RNA metabolism, including transcription, splicing, transport, and stability (Zeng et al., 2024). Under physiological conditions, TDP-43 predominantly resides in the nucleus but shuttles between the nucleus and cytoplasm. In pathological states, TDP-43 becomes mislocalized to the cytoplasm, where it forms insoluble, ubiquitinated aggregates, a defining feature observed in approximately 97% of ALS and 45% of FTD cases (Ling et al., 2013; Neumann et al., 2006). Mutations in the TARDBP gene, encoding TDP-43, have been identified in familial forms of ALS and FTD, further implicating TDP-43 in the pathogenesis of these diseases (Sreedharan et al., 2008).

Synaptic dysfunction and loss are early events in the progression of NDs and are closely associated with cognitive decline and behavioral deficits. In ALS and FTD, synaptic alterations have been documented in both cortical and hippocampal regions, correlating with impairments in memory, executive function, and social behavior (Mora and Allodi, 2023). Experimental models have demonstrated that TDP-43 loss of function leads to synaptic deficits, including reduced dendritic spine density and altered synaptic protein expression, contributing to neuronal dysfunction (Ni et al., 2023). Mitochondrial abnormalities further exacerbate synaptic dysfunction, as pathological TDP-43 disrupts mitochondrial dynamics, trafficking, and oxidative phosphorylation, contributing to bioenergetic failure and oxidative stress (Gao et al., 2019).

In this study, we employed an inducible mouse model expressing a cytoplasmically mislocalized form of human TDP-43 lacking a functional nuclear localization signal (ΔNLS). This model recapitulates key features of TDP-43 proteinopathy, including cytoplasmic aggregation, altered gene expression, and behavioral impairments such as motor abnormalities, reduced sociability and working memory deficits (Alfieri et al., 2014; Igaz et al., 2011). These phenotypes appeared at early time points (0.5-1 month) after TDP-43-ΔNLS induction. Previous investigations have reported time-dependent neurodegeneration in these mice, with only mild neuronal loss at 1 month post transgene induction versus more pronounced cell death after 6 months of TDP-43-ΔNLS expression (Alfieri et al., 2014). To further elucidate the early structural correlates of the observed behavioral phenotypes, we conducted ultrastructural analyses of cortical and hippocampal regions, focusing on synaptic and mitochondrial integrity.

## 2. Materials and Methods

### 2.1 Animals

We used our previously generated and characterized CamK2/TDP-43-ΔNLS mouse model for both immunofluorescence and TEM analysis. tetO-TDP-NLS4 line was generated as described in Igaz et al. (2011). Monogenic tetO-TDP-NLS4 mice were bred to CamK2a-tTA mice producing non-transgenic mice (nTg), tTA monogenic, monogenic tetO-TDP-NLS4 (non TDP-43-ΔNLS expressing mice) and bigenic mice expressing TDP-43-ΔNLS (referred hereafter as ΔNLS). Breeding animals were treated with 0.2 mg/ml doxycycline hyclate (Dox; sc-204734A; Santa Cruz Biotechnology) in drinking water to avoid pre- and postnatal developmental effects due to the transgene expression (Tg) in pups. At weaning (post-natal day 28) pups were switched to regular water to allow Tg expression and mice were analyzed at 1-month (1mo, 60 postnatal days) Tg induction. Mice from both sexes were used, as in our previous studies (Alfieri et al., 2014; Alfieri et al., 2016; Charif et al., 2020; Igaz et al., 2011; Silva et al., 2019). From ear biopsies, genomic DNA was extracted to screen for presence of the Tg by PCR amplification with the primers TDP-forward (TTGGTAATAGCAGAGGGGGTGGAG), MoPrP-reverse (TCCCCCAGCCTAGACCACGAGAAT), Camk2a-tTA-forward (CGCTGTGGGGCATTTTACTTTAG), and Camk2atTA-reverse (CATGTCCAGATCGAAATCGTC). (Alfieri et al., 2014; Igaz et al., 2011).

The experimental protocol for this study was approved by the National Animal Care and Use Committee (CICUAL) of the University of Buenos Aires.

### 2.2 Transmission Electron microscopy (TEM)

#### 2.2.1 Brain tissue collection

Animals were deeply anesthetized with an intraperitoneal injection of pentobarbital (0,2 mg/g) followed by transcardial perfusion with ice-cold 0.1 M Phosphate Buffer Saline (PBS) pH=7.4 and 10u/ml heparine. The brains were rapidly extracted and thick brains sections (approximately Bregma -2.80 mm) were dissected in 1mm^3^ cubes corresponding to Hippocampus (Hp) and Cortex (Ctx), then postfixed in 2.5% glutaraldehyde in 0.1 M phosphate buffer, pH 7.4 for 4 hours at 4°C. This was followed by two washes with 0.1 M phosphate buffer, pH 7.4.

#### 2.2.2 TEM staining

Dissected samples of 1 mm^3^ corresponding to CA1 area of Hp and somatosensory Ctx were post-fixed in 1% (w/v) osmium tetroxide in 0.1 M phosphate buffer for 30 min. After dehydration in ethanol gradient, tissues were contrasted with 5% (w/v) uranyl acetate and then embedded in Durcupan (Fluka AG, Chemische Fabrik, Buchs SG, Switzerland). After sections were embedded, 1-μm slices were obtained and toluidine blue stained, to select similar areas of CA1 and Ctx. Then ultrathin sections (50 nm) were obtained and stained with lead citrate (Reynolds, 1963), as previously described (Soriano et al., 2020).

#### 2.2.3 Image acquisition

Images were acquired on a Zeiss 109 TEM (Carl Zeiss, Oberkochen, Germany) and photographed with a GATAN CCD camera (Pleasanton, CA, USA).

#### 2.3.4 Synaptic and mitochondrial quantitative analysis

For the quantification of synapses and mitochondria, electron microscopy images at 20000x magnification were used. The number of synapses per 100 μm^2^ was measured in CA1 area of the Hp and in the Ctx using imageJ (version 1.5f). Synapses were identified by the presence of two opposing membranes and synaptic vesicles. A similar analysis was performed to quantify the number of mitochondria per 100 μm^2^. Mitochondria were identified based on the presence of double membranes and cristae. To assess synapse morphology, electron microscopy images at 50000x magnification were used. Synapse length was defined as a straight line from one end of the synaptic contact to the other. Synaptic cleft width was measured as the distance between the opposing membranes, and the thickness of the postsynaptic density (PSD) was determined as the width of the electron-dense region on the postsynaptic side of the synapse. All measurements were performed using ImageJ.

### 2.4 Analysis of transcriptomic and proteomic datasets

To explore whether the ultrastructural alterations observed in CamK2-TDP-43-ΔNLS mice were accompanied by early molecular changes, transcriptomic and proteomic datasets from previously published studies on ΔNLS mouse models of TDP-43 proteinopathy (Igaz et al., 2011; San Gil et al., 2024) were reanalyzed. We focused specifically on genes related to synaptic structure and function, using annotations from the Gene Ontology categories “regulation of synapse structure or activity” (GO:0050803) and “synapse organization” (GO:0050808) that were identified as significantly altered by San Gil et al. (2024). A curated list of genes from these categories was compiled and examined for differential expression at early disease stages. Data analysis and visualization were carried out in the R environment. Log□ fold changes (LFC) and corresponding p-values were used to generate volcano plots, which highlighted genes with significant expression changes and functional relevance to synaptic biology. The datasets analyzed included results from the CamK2-TDP-43-ΔNLS mice (Igaz et al., 2011) with forebrain neuron expression, and the rNLS8 transgenic mouse model, which expresses the same TDP-43-ΔNLS construct under a pan-neuronal promoter (Walker et al., 2015). Additional proteomic data from rNLS8 mice (San Gil et al., 2024) and transcriptomic and proteomic human ALS and FTD tissue (Hasan et al., 2022; Umoh et al., 2018) were retrieved from the public webtool “TDP-MAP” (https://shiny.rcc.uq.edu.au/TDP-map/).

### 2.5 Statistical analysis

Data from TEM analysis of different synaptic and mitochondrial measurements are expressed as mean ± SEM and statistical analysis was performed using Prism 8 (GraphPad Software). Student’s t-test was used to compare control and ΔNLS groups. Statistical significance was defined as *p* <0.05.

## 3. Results

Previous studies have shown that alterations in TDP-43 expression levels, subcellular location or mutations are capable of causing changes in the ultrastructure of different brain cell types, including neurons and glia (Davis et al., 2018; Jara et al., 2019; Wang et al., 2019). Since TDP-43-ΔNLS mice show progressive neurodegeneration (Alfieri et al., 2014; Igaz et al., 2011), we chose to analyze a time point with limited neuronal loss but with florid presentation of behavioral phenotypes, including motor, cognitive and social deficits (Alfieri et al., 2014). An ideal, early time point after transgene induction that presents these features is 1 month after Dox removal (1mo); therefore, our study was performed entirely in this time period, representing an early phase of the disease progression.

### 3.1 Cytoplasmic TDP-43 expression in forebrain neurons leads to multiple ultrastructural abnormalities in mouse brain regions

An initial qualitative assessment using TEM analysis of cortical and hippocampal tissue of TDP-43-ΔNLS mice revealed multiple ultrastructural abnormalities. We focused on features involving both intra and extracellular structures in different cell types.

We found clear evidence of altered ultrastructural features in the brains of TDP-43-ΔNLS mice, including altered axons with neurofilament disorganization (**Fig. 1A-B**), perivascular edema (**Fig. 1C-D**), and edema within astrocytes (**Fig. 1E-F**). These anomalies were found in both hippocampal and cortical regions of transgenic mice. Further pathological features found in the brains of TDP-43-ΔNLS mice include edema and enlarged lumen of rough endoplasmic reticulum (**Fig. 1G-H**), abnormal mitochondria (**Fig. 1I-J**) and neurons with abnormal morphology, intracellular edema and condensed nuclei (**Fig. 1K-L**), both in the hippocampus (**Fig. 1G, I, K**) and the cortex (**Fig. 1H, J, L**) of mice conditionally expressing cytoplasmic TDP-43.

**Figure 1.**
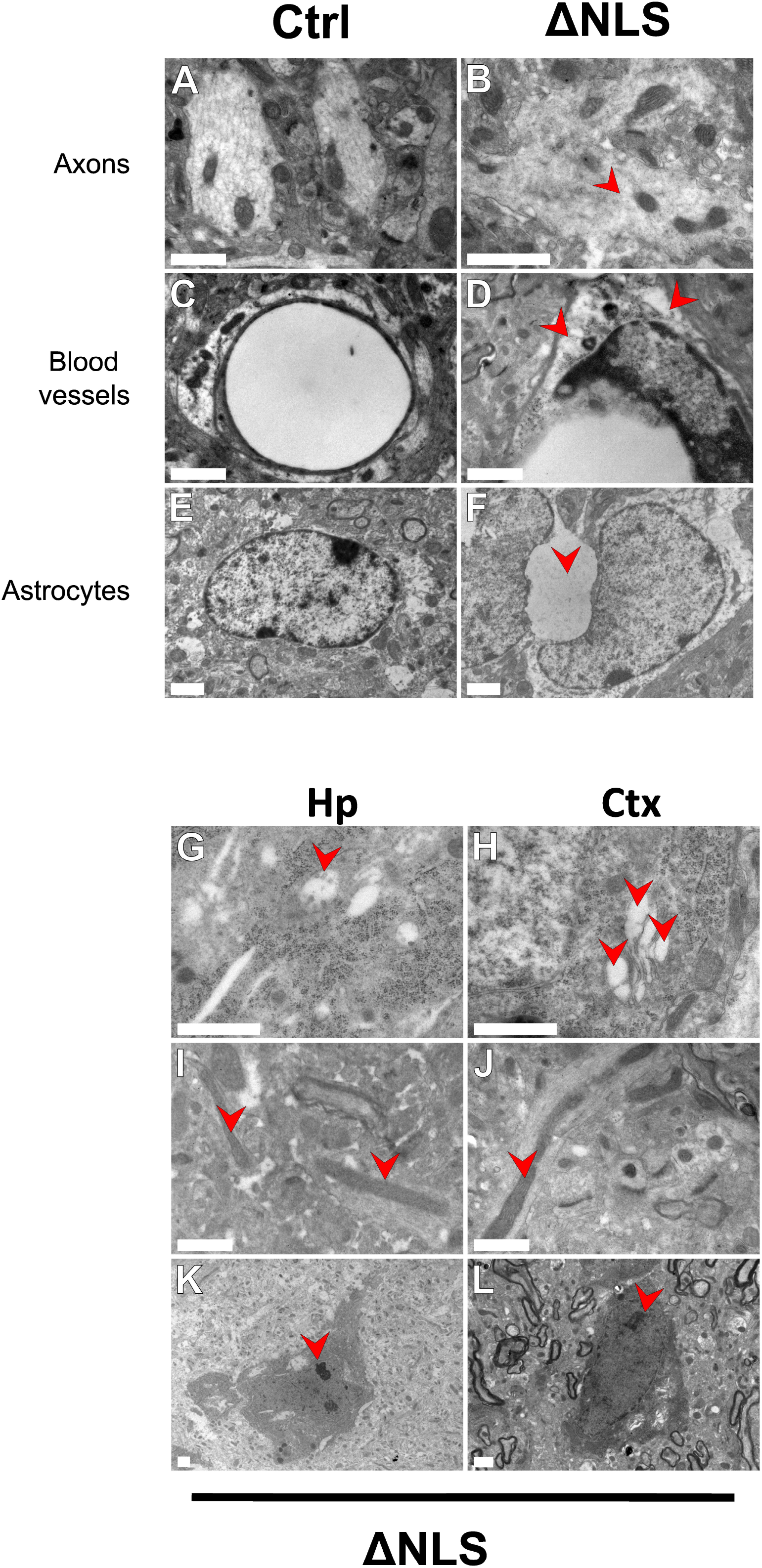
Ultrastructural abnormalities in the brains of TDP-43-ΔNLS mice. **(A–B)** Representative transmission electron microscopy (TEM) images of axons. **(A)** Normal axon morphology in a control (Ctrl) mouse. **(B)** Pathological axon in a TDP-43-ΔNLS (ΔNLS) mouse showing disorganized microtubules (red arrowhead). **(C–D)** Micrographs of blood vessels. **(C)** Normal vascular structure in Ctrl mice. **(D)** Blood vessels from a ΔNLS mouse exhibiting signs of edema (red arrowheads). **(E–F)** Astrocyte morphology. **(E)** Normal astrocyte in a Ctrl mouse. **(F)** Astrocyte from a ΔNLS mouse showing cellular edema (red arrowhead). **(G-L)** Further examples of ultrastructural abnormalities in the hippocampus (Hp) and cortex (Ctx) of ΔNLS mice. **(G-H)** Red arrowheads highlight edema and enlarged lumen of rough endoplasmic reticulum. (I-J) Green arrowheads highlight abnormal mitochondria. **(K-L)** The blue arrowheads indicate neurons with abnormal morphology, intracellular edema, and condensed nuclei. Scale bar = 1 μm.

### 3.2 TDP-43-ΔNLS mice display reduced cortical and hippocampal synaptic density

A common feature of multiple NDs is the alteration of synapse number and morphology (Meftah and Gan, 2023; Uytterhoeven et al., 2025). In order to assess if the short-term expression of TDP-43-ΔNLS led to changes in synaptic density, we used TEM analysis in Ctx and Hp, two disease-relevant regions of the brain (**Fig. 2A**). Cortical synaptic density was significantly decreased by 43.5 % (25.47 ± 1.86 vs. 14.38 ± 1.07 synapses/100 □ m^2^ in Control vs. TDP-43-ΔNLS mice, respectively; p < 0.0001) (**Fig. 2B, left**). We also found a similar reduction in synaptic density in the Hp, significantly decreasing by 36.2 % (27.66 ± 2.03 vs. 17.66 ± 1.34 synapses/100 □ m^2^ in Control vs. TDP-43-ΔNLS mice, respectively; p = 0.0001) (**Fig. 2B, right**). This remarkable early synapse loss is occurring at a time point where multiple behavioral domains, depending on the proper function of these brain regions, are severely affected (Alfieri et al, 2014).

**Figure 2.**
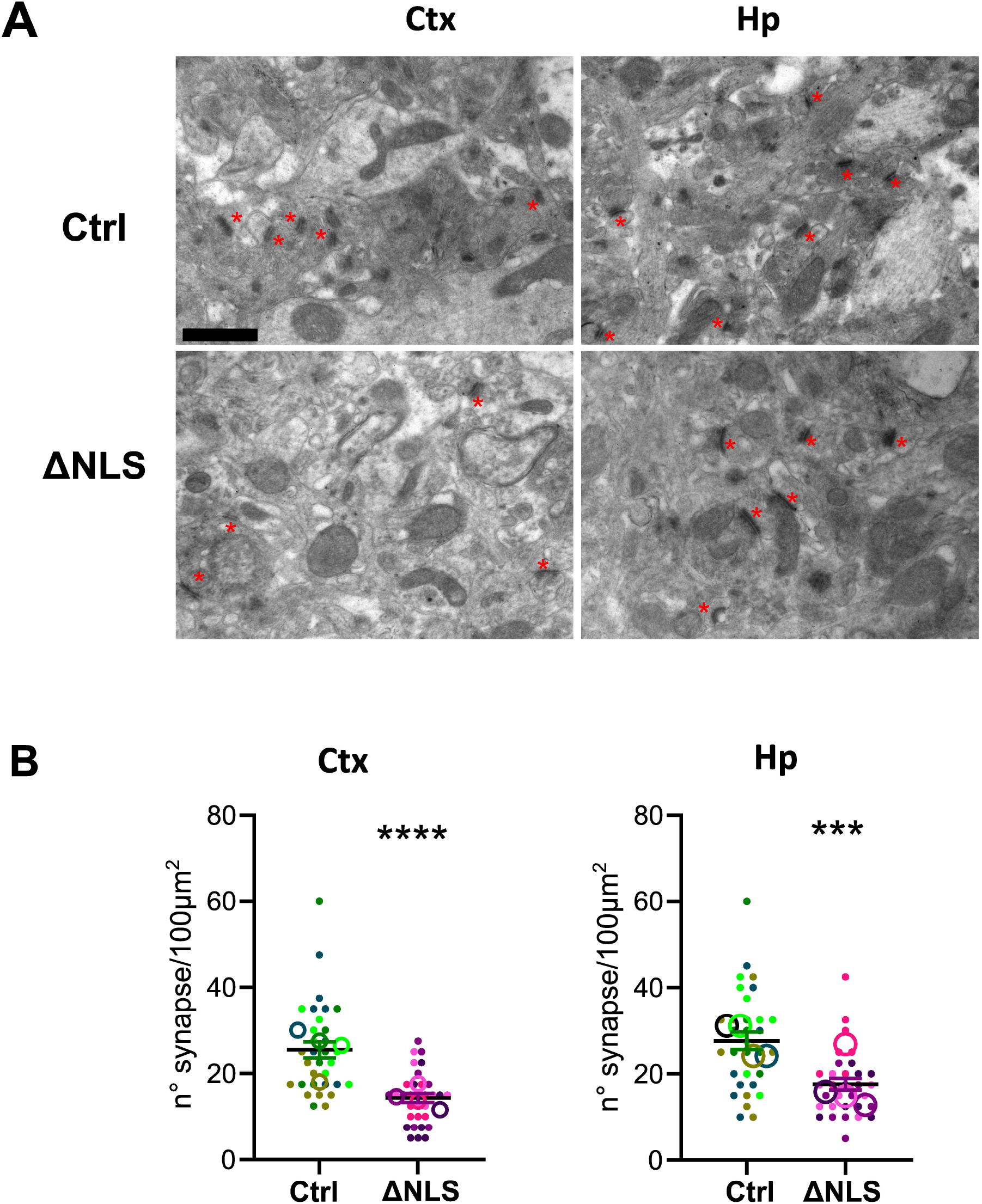
Reduced synaptic density in the cortex and hippocampus of TDP-43-ΔNLS mice. **(A)** Representative TEM images of the Ctx and Hp from control (Ctrl) and TDP-43-ΔNLS (ΔNLS) mice. Red asterisks indicate synapses (*). **(B)** Quantification of synapse number per 100 μm^2^ in Ctx and Hp. ΔNLS mice exhibit significantly reduced synaptic density compared to controls. Colored dots represent synapse counts from different fields within individual animals; large circles indicate the average value for each animal. Data are presented as mean ± SEM. ***p < 0.001, ****p < 0.0001 compared to Ctrl (Student’s t-test), n = 8 fields per animal, 4 animals per group. Scale bar = 1 μm.

### 3.3 Expression of TDP-43-ΔNLS alters the fine structure of cortical and hippocampal synapses

To further characterize this synaptic phenotype, we set out to assess if the expression of cytoplasmic TDP-43 is affecting not only synapse number and/or density but other subtler morphological synaptic features (**Fig. 3**). Higher magnification TEM images show more detailed characteristics of cortical and hippocampal synapse structure in Control and transgenic mice (**Fig 3A-D** and their insets). Since there is recent evidence for a direct relationship between synapse size and strength (Holler et al., 2021), we quantified synapse length in transgenic and control brains, and found a significant increase both for cortical (**Fig. 3E**) and hippocampal synapses (**Fig. 3H**). In the Ctx, synapse length was 26.0 % higher in ΔNLS than controls (Control: 202.8 ± 8.5 nm vs.TDP-43-ΔNLS 255.6 ± 11.6 nm, p = 0.0002), while in the Hp it was increased by 31.9 % (Control: 183.0 ± 7.4 nm vs.TDP-43-ΔNLS 241.4 ± 11.0 nm, p < 0.0001).

**Figure 3.**
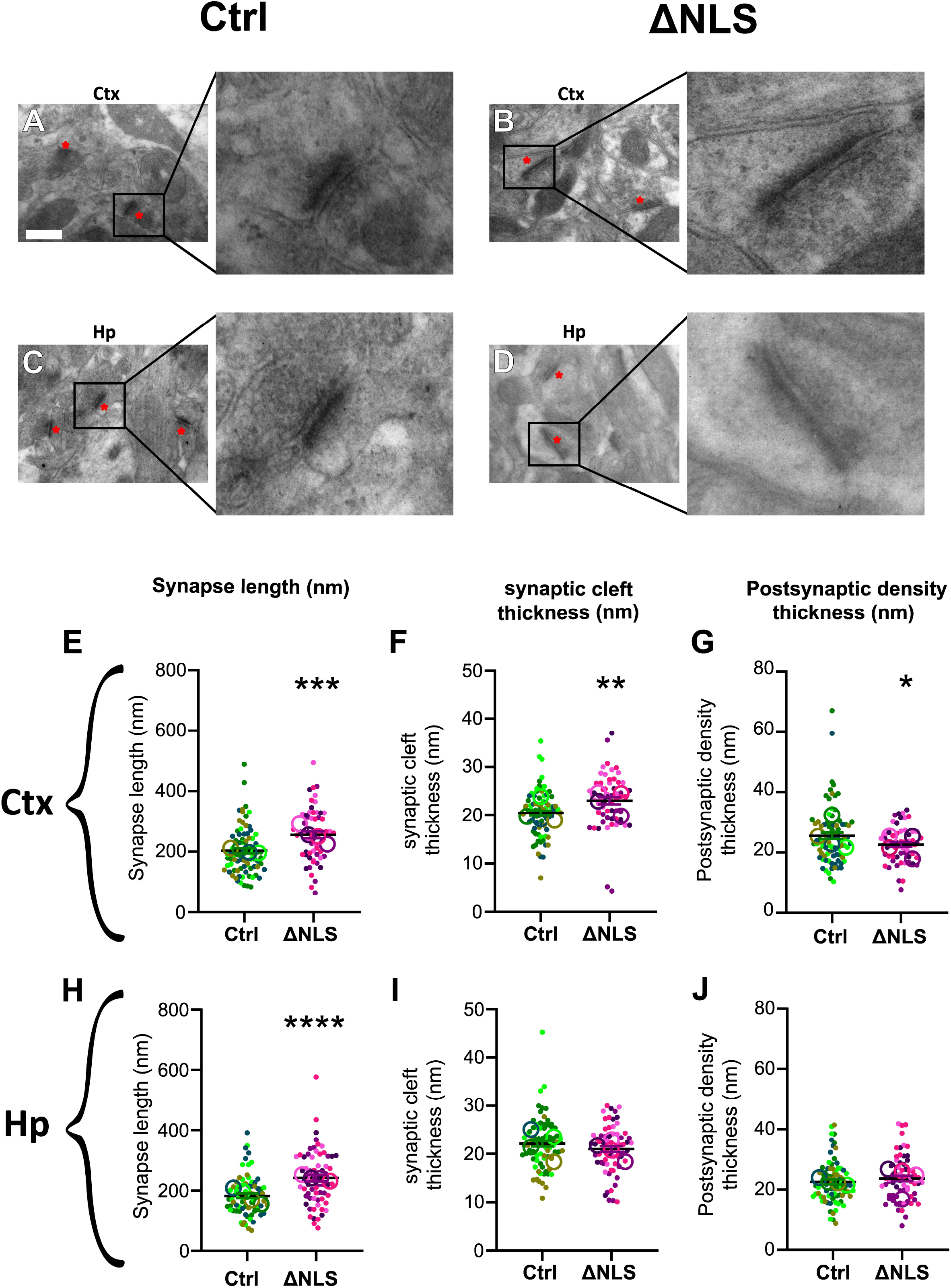
TDP-43-ΔNLS expression alters specific ultrastructural features of synapses. Representative TEM images of synapses of Ctrl **(A**,**C)** and ΔNLS **(B**,**D)** for Ctx and Hp respectively. Synapses are marked with black asterisks. Insets show magnified views highlighting the presynaptic and postsynaptic membranes, synaptic cleft, and postsynaptic density (PSD). **(E–J)** Quantification of synaptic parameters: synapse length **(E, H)**, synaptic cleft width **(F, I)**, and PSD thickness **(G, J)** in Ctx and Hp. Measurements were taken from 50,000X magnification images. Colored dots represent individual synapses; circles represent the average value for each animal. Data are shown as mean ± SEM. *p < 0.05, **p < 0.01, ***p < 0.001, ****p < 0.0001 versus Ctrl (Student’s t-test), n = 8 fields per animal, 4 animals per group. Scale bar = 400 nm.

We also examined the synaptic cleft width, as synaptic cleft geometry is an important factor in shaping synaptic currents (Savtchenko and Rusakov, 2007). Across many synaptic types, this distance lies within a relatively narrow range, likely determined by a rigid intracleft protein scaffolding structure (Zuber et al., 2005). Synapses in the Ctx (**Fig. 3F**) of transgenic mice showed a 12.5 % increase in synaptic cleft width (Control: 20.47 ± 0.51 nm vs.TDP-43-ΔNLS 23.03 ± 0.74 nm, p = 0.0041); however, in hippocampal synapses (**Fig. 3I**) did not differ significantly (Control: 22.12 ± 0.60 nm vs.TDP-43-ΔNLS 20.99 ± 0.59 nm, p = 0.1865).

Finally, we evaluated the thickness of the PSD. It is generally accepted that there is a complex relationship between PSD size and synaptic strength (Dosemeci et al., 2016). Cortical PSD thickness (**Fig. 3G**) was significantly decreased in TDP-43-ΔNLS mice by 11.8 % (Control: 24.70 ± 1.05 nm vs.TDP-43-ΔNLS 21.79 ± 0.77 nm, p = 0.042), while this parameter was not changed in the Hp (**Fig. 3J**) (Control: 22.54 ± 0.81 nm vs.TDP-43-ΔNLS 23.65 ± 0.89 nm, p = 0.3581).

Overall, these data show that a complex rearrangement of synaptic morphology takes place in the brain of mice expressing cytoplasmic TDP-43, most notably in cortical regions.

### 3.4 TDP-43-ΔNLS mice show region-specific differences in brain mitochondrial density

In addition to synaptic pathology, abnormalities in mitochondrial morphology and/or function have been reported in both animal models and human ND (Briston and Hicks, 2018). Moreover, TDP-43 has been linked to mitochondrial physiology and pathology (Jiang and Ngo, 2022; Shan et al., 2010; Stribl et al., 2014; Wang et al., 2016; Xu et al., 2010). Thus, we first investigated if overexpression of cytoplasmic TDP-43 has an impact on mitochondrial density in the brain (**Fig. 4**). Surprisingly, we found a bidirectional regulation in different brain regions of TDP-43-ΔNLS mice (**Fig. 4A-B**), as the Ctx showed a 28 % increase (21.72 ± 1.16 vs. 27.81 ± 1.46 mitochondria/100 □ m^2^ in Control vs. TDP-43-ΔNLS mice, respectively; p < 0.0018) while the Hp displayed a 22.1 % reduction (29.75 ± 2.15 vs. 23.19 ± 1.70 mitochondria/100 □ m^2^ in Control vs. TDP-43-ΔNLS mice, respectively; p < 0.0198).

**Figure 4.**
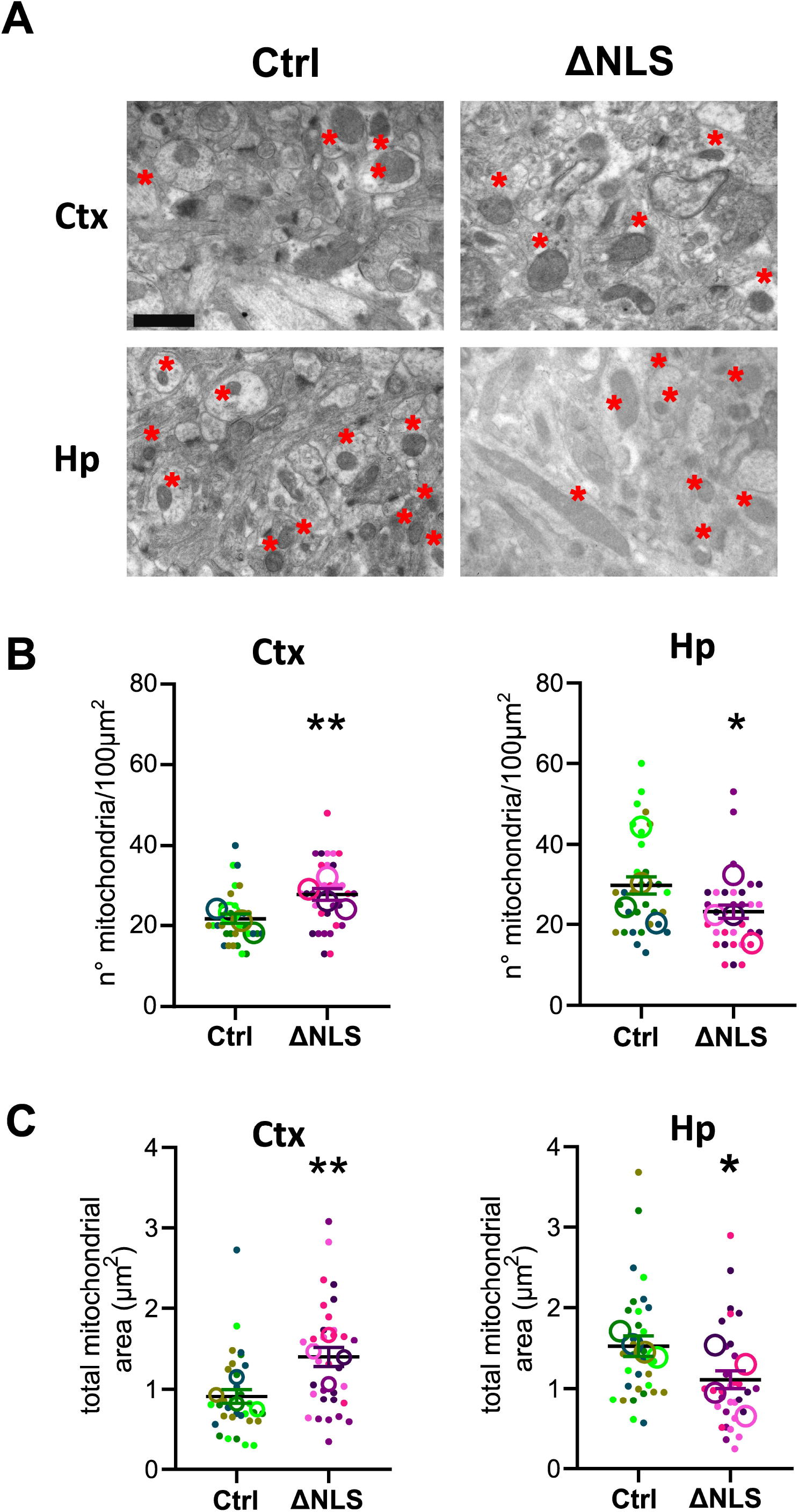
TDP-43-ΔNLS mice show region-specific changes in mitochondrial density and total area. **(A)** Representative TEM images of mitochondria (red asterisks) in the Ctx and Hp of Ctrl and ΔNLS mice. **(B)** Mitochondrial density (number per 100 μm^2^) in Ctx and Hp. ΔNLS mice show increased mitochondrial density in the Ctx, while a significant decrease is observed in the Hp. **(C)** Total mitochondrial area per region, calculated as the sum of individual mitochondrial areas. An increase in total area is observed in Ctx and a decrease in Hp in ΔNLS mice compared to Ctrl. Colored dots represent individual field values; circles indicate the animal average. Data are shown as mean ± SEM. *p < 0.05, **p < 0.01 versus Ctrl (Student’s t-test), n = 8 fields per animal, 4 animals per group. Scale bar = 1 μm.

To study if this change in mitochondrial density reflected a change in mitochondrial size, we first analyzed the total area covered by mitochondria (**Fig. 4C**). Cortical samples presented a statistically significant 53.8 % increase (Control: 0.91 ± 0.09 mitochondria/ □ m^2^ vs.TDP-43-ΔNLS 1.40 ± 0.12 mitochondria/ □ m^2^, p = 0.0016), while this parameter was significantly decreased by 27.5 % in the Hp (Control: 1.53 ± 0.13 mitochondria/ □ m^2^ vs.TDP-43-ΔNLS 1.11 ± 0.11 mitochondria/ □ m^2^, p = 0.0158). This correlation of direction of change between mitochondrial density and total mitochondrial area suggested that these region-specific differences were not mainly caused by altered mitochondrial dynamics (i.e. fission or fusion), but rather due to a change in the total mitochondrial number.

### 3.5 Additional features of mitochondrial changes in TDP-43-ΔNLS mouse brain regions

In order to examine in more detail the nature of the effect of abnormal TDP-43 localization on mitochondria morphology, we quantified features of individual mitochondria (**Fig. 5**) instead of averaged fields, as in **Fig 4**. Individual area of mitochondria showed no significant differences in either Ctx (Control: 0.105 ± 0.006 □ m^2^ vs.TDP-43-ΔNLS 0.119 ± 0.007 □ m^2^, p = 0.1057) or Hp (Control: 0.127 ± 0.008 □ m^2^ vs.TDP-43-ΔNLS 0.120 ± 0.009 □ m^2^, p = 0.5801) (**Fig 5A**). Frequency distribution analysis of individual mitochondrial size in both areas revealed no differences in relative frequency distribution (**Fig. 5B**). This result is in line with our interpretation that changes in total mitochondrial area is mainly due to a change in total mitochondrial number and not in individual size.

**Figure 5.**
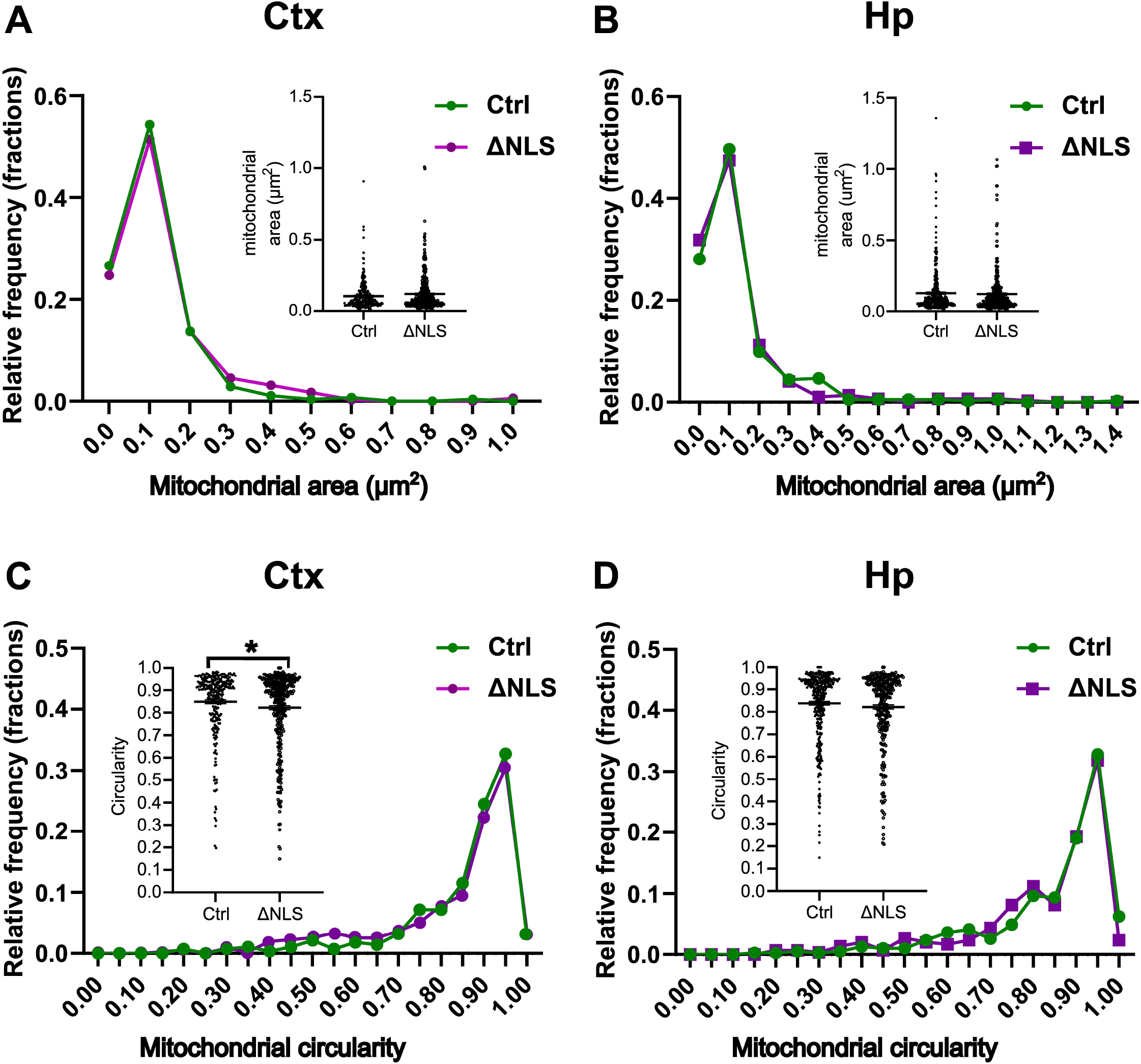
Comparable mitochondrial size distribution but altered circularity in the cortex of TDP-43-ΔNLS mice. **(A–B)** Relative frequency distribution of individual mitochondrial area in Ctx **(A)** and Hp **(B)**. Insets show raw area measurements per mitochondrion. **(C–D)** Relative frequency distribution of mitochondrial circularity in Ctx **(C)** and Hp **(D)**. Insets display individual circularity values. Circularity is significantly reduced in Ctx of ΔNLS mice, while no significant differences are observed in Hp. Data are shown as mean ± SEM. *p < 0.05 versus Ctrl (Student’s t-test), n = 8 fields per animal, 4 animals per group. Scale bar = 400 nm.

Finally, we measured mitochondrial circularity, to gauge more subtle morphological abnormalities in mitochondrial structure. Cortical mitochondria (**Fig. 5C**) showed a small (4 %) but significant decrease in circularity (Control: 0.849 ± 0.009 vs.TDP-43-ΔNLS 0.822 ± 0.009, p = 0.0379). Relative frequency analysis suggested that this decrease was due to a relative increase in mitochondria with low (0.4-0.6) circularity (i.e. much more elongated; see **Fig. 1I-J**). On the other hand, mitochondria from the Hp (**Fig. 5D**) showed no significant changes in circularity (Control: 0.839 ± 0.008 vs.TDP-43-ΔNLS 0.821 ± 0.010, p = 0.1786).

In sum, analysis of mitochondrial ultrastructural features showed a bidirectional regulation in cortical vs hippocampal areas: the former displaying increased mitochondrial density and total covered area, while the latter presenting decreased mitochondrial density and total covered area. Both regions showed no changes in individual mitochondrial area, but the cortical mitochondria additionally exhibited significantly decreased circularity.

### 3.6 Transcriptomic/proteomic analysis from transgenic TDP-43-ΔNLS mice and human proteinopathy show widespread decreased expression of synaptic structure and function genes

Our electron microscopy findings point to an alteration of synaptic number and structure in the CamK2-TDP-43-ΔNLS mouse model of TDP-43 proteinopathy. To investigate if these early changes are mirrored in transcriptomic and proteomic changes in both TDP-43-ΔNLS mouse models and FTD human tissue, we re-analyzed 1) our microarray data from early-stage CamK2-TDP-43-ΔNLS mice (Igaz et al., 2011) and 2) published datasets from previous sources using the webtool “TDP-MAP” (https://shiny.rcc.uq.edu.au/TDP-map/) (San Gil et al., 2024), which allows targeted meta-analysis of available datasets (**Fig. 6**). We focused on genes belonging to two Gene Ontology (GO) categories revealed to be significantly changed in a cortical proteomic analysis of the rNLS8 mouse model (San Gil et al., 2024), which is generated by crossing the same TDP-ΔNLS protein as in our mice, but driven by the pan-neuronal promoter of the NEFH gene (Walker et al., 2015). The categories selected were GO:0050803 (regulation of synapse structure or activity) and GO:0050808 (synapse organization), which we considered relevant for our ultrastructural synaptic analysis. Re-analysis of genes in these GO categories showed that, in CamK2-TDP-43-ΔNLS 2 weeks after induction, cortical mRNAs coding for most of these proteins were significantly decreased (**Fig. 6A-B**). When analyzing rNLS8 mouse gene expression, we found that after 1, 2, 4 or 6 weeks post transgene induction or a “recovery” situation after transgene suppression for 2 additional weeks, the same pattern of decrease was apparent at every time point, indicating widespread reduction of synaptic structure/organization-related proteins (**Fig. 6C,E**).

**Figure 6.**
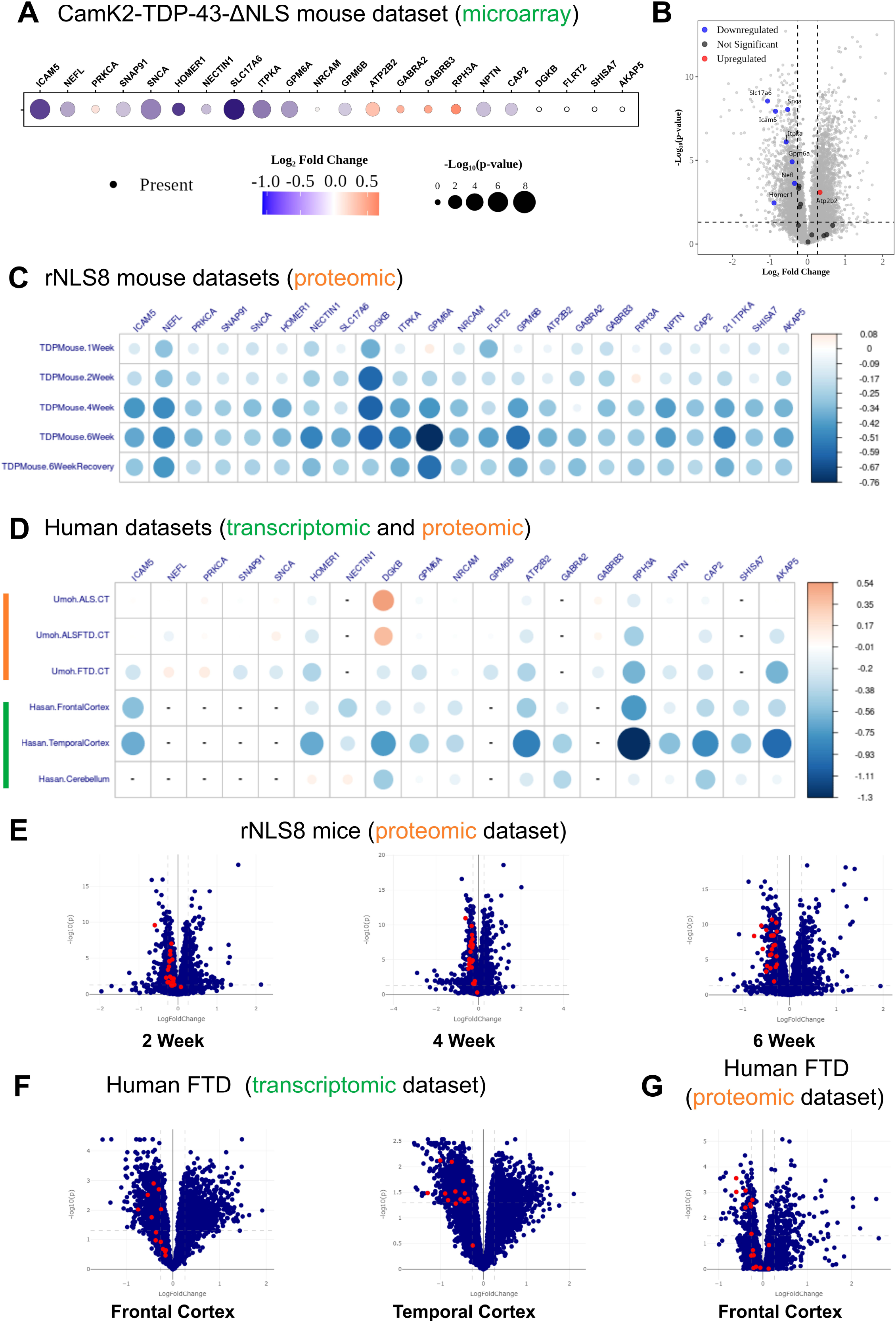
Transcriptomic and proteomic analyses reveal broad downregulation of synaptic genes in TDP-43-NLS mice and human TDP-43 proteinopathy. **(A)** Bubble plot of synaptic genes mRNA abundance in microarray data from the Ctx of CamK2-TDP-43-ΔNLS mice (modified from Igaz et al. (2011). **(B)** Volcano plot of transcript mean log fold change (LFC) and significance level [-log10(Pvalue)] from the cortex of CamK2-TDP-43-ΔNLS mice, showing highlighted transcripts from the Gene Ontology categories “regulation of synapse structure or activity” (GO:0050803) and “synapse organization” (GO:0050808). Transcripts with no significant change (grey), significantly upregulated (red) or downregulated (blue) are shown. **(C)** Plot of synaptic genes/protein abundance in proteomic data from the cortex of rNLS8 mice (San Gil et al., 2024) (top panel) or transcriptomic (Hasan et al., 2022) and proteomic (Umoh et al., 2018) of human post-mortem tissue (bottom panel) Bubble plots were custom-generated using R **(A)** or generated with the TDP-map webtool **(C-D)** and data points were color-scaled to illustrate the LFC where red is increased and blue is decreased compared to controls. Circle size is dependent on the magnitude of LFC, the larger the change the larger the circle size. **(E)** Volcano plots of cortical protein level mean LFC (rNLS8/Ctrl) and significance level [-log10(Pvalue)] from mice at onset (2 wk) (left panel), early (4 wk) (middle panel) and late (6 wk) (right panel) disease. **(F)** Volcano plots of transcript mean LFC and significance level [-log10(Pvalue)] of human FTD frontal **(F)** and temporal **(G)** Ctx (Hasan et al., 2022). **(G)** Volcano plot of protein level mean LFC and significance level [-log10(Pvalue)] of human FTD frontal Ctx (Umoh et al., 2018). The red dots in the volcano plots represent the transcripts/proteins from the Gene Ontology categories “regulation of synapse structure or activity” (GO:0050803) and “synapse organization” (GO:0050808).

To investigate if these alterations occurring in both early and late stage disease of TDP- 43- ΔNLS mouse model reflects changes seen in end-stage human TDP-43 proteinopathy tissues, we performed a similar analysis in two recent human datasets: I) proteomics from post-mortem frontal cortical tissues from clinically defined (sporadic) ALS, ALS/FTLD, FTLD-TDP and control cases (Umoh et al., 2018), and II) transcriptomics from post-mortem frontal cortex, temporal cortex, and cerebellum from FTLD-TDP and control cases (Hasan et al., 2022). Remarkably, most synaptic structure/organization proteins that were decreased in the CamK2-TDP-43-ΔNLS and in the rNLS8 mouse cortex in disease and recovery stages were also significantly decreased in human ALS and FTD post-mortem brain tissues (**Fig. 6D and 6F-G**). These data indicate that human disease-relevant synaptic proteins are persistently decreased in both mouse models expressing cytoplasmic TDP-43, and that these findings recapitulate changes found in late-stage human TDP-43 proteinopathies.

## 4. Discussion

Previous work from our group has contributed to the characterization of the CamK2-TDP-43-ΔNLS mouse model through behavioral assessments, neurodegeneration analysis, and protein expression profiling. This inducible model, which conditionally overexpresses cytoplasmically mislocalized TDP-43, recapitulates key pathological features of FTD and ALS, including altered social behavior, working memory deficits, spasticity, and motor coordination impairments (Alfieri et al., 2014; Alfieri et al., 2016; Igaz et al., 2011). To bridge these phenotypes with potential underlying mechanisms, we studied ultrastructural synaptic and features in the Cx and Hp of this animal model of TDP-43 proteinopathies.

Synaptic integrity is affected in neurodegenerative diseases such as ALS and FTD (Wilson et al., 2023). TEM analysis performed in postmortem tissue of ALS patients evidenced a significant loss of synapses in the prefrontal cortex (Henstridge et al., 2018). A similar finding occurred in bvFTD (behavioral variant FTD) patients, showing synaptic loss in the frontotemporal region compared to control patients in PET studies (Malpetti et al., 2023). Moreover, PET imaging using the synaptic vesicle protein tracer 18F-SynVesT-1 revealed decreased synaptic density in the frontal and temporal lobes of ALS patients (Tang et al., 2022). Such ultrastructural changes can significantly impact synaptic function; thus, morphological alterations in synapses will affect the transmission of action potentials, as well as learning and memory. In aging synapses, the reduced number of synaptic vesicles and mitochondria may disrupt normal neurotransmitter release, leading to impaired neural signal transduction (Fan et al., 2018).

Until now, no other studies have analyzed the ultrastructural features of the brains of TDP-43-ΔNLS mice. We aimed to explore whether the behavioral alterations displayed by our model had a basis in ultrastructural changes in the Cx and Hp. We demonstrate that early overexpression of TDP-43-ΔNLS leads to a reduction in synaptic density in both cortical and hippocampal regions. Similar synaptic alterations have been described in other TDP-43 transgenic models such as in the TDP-43^A315T^ mice that exhibit reduction of dendritic spine density prior to neurodegeneration (Handley et al., 2017). Interestingly, *in vivo* microglial TDP-43 depletion led to synapse loss in a transgenic mouse model of AD (Paolicelli et al., 2017). Also, neuronal overexpression of hTDP-43 in mice provoked synaptic loss, evidenced by reduced synaptophysin levels (Medina et al., 2014). Moreover, overexpression of a mutant form of hTDP-43 —associated with ALS— in a mouse model of proteinopathies resulted in structural alterations of synapses in cortical pyramidal neurons. The expression of TDP-43^A315T^ was mainly nuclear but cytoplasmic localization was also observed. These alterations may be driving the reduced excitability observed (Jiang et al., 2019). This phenomenon was also reported in other NDs, as electron microscopy analysis of cortical samples from patients with AD showed a decrease in synaptic density in the neuropil of both Layers II and III (Dominguez-Alvaro et al., 2021). Collectively these findings support the idea that synaptic dysfunction likely precedes overt neuronal loss.

Notably, cortical synapses of TDP-43-ΔNLS transgenic mice exhibited ultrastructural alterations characterized by increased synaptic cleft width and a significant thinning of the PSD. In the Hp, only the synapse length was significantly greater than control. Since PSD thickness and the synaptic cleft width are related with synaptic transmission efficiency and plasticity (Di and Zheng, 2013), the thinning of the PSD and the increased synaptic cleft width may reflect defective synaptic transmission which could contribute to the behavioral deficits described previously in our model (Alfieri et al., 2014). Consistent with these findings, pathway analysis of significantly changing genes between bigenic and control mice described a downregulation of synaptic transmission-related genes in the Ctx of TDP-43-ΔNLS mice 10 days post induction, as revealed by RNAseq analysis (Amlie-Wolf et al., 2015). Recently, a study using induced pluripotent stem cell-derived cortical neurons from FTD patients demonstrated reduced number and altered morphologies of dendritic spines and significantly altered synaptic function (Huber et al., 2025). Lastly, a very recent report showed that TDP-43 reduction in human stem-cell derived neurons induces cryptic splicing and downregulation of synaptic and membrane excitability genes, whose misregulation drives neuronal dysfunction. These cryptic splicing events have been found in postmortem brains from FTD patients, highlighting the potential role of synaptic deficits in human TDP-43 proteinopathies (Guo et al., 2025).

Dysfunction in mitochondria is a common sign present in many NDs such as AD, Huntington’s and Parkinson’s disease (Martin et al., 2015; Ryan et al., 2015). Given the high energy demands of synaptic maintenance and the critical role of mitochondria— particularly their enrichment in synaptic boutons—in supporting synaptic function (Li and Sheng, 2022), we sought to analyze mitochondrial morphology and abundance in our model. In the Hp, ΔNLS mice showed reduced mitochondrial density and total mitochondrial area. Conversely, in the Ctx, both mitochondrial density and total area were increased compared to controls. Importantly, mitochondrial size remained unchanged in both regions, probably suggesting that mitochondrial dynamics, specifically the balance between fission and fusion processes, are not significantly affected at this stage. TEM studies in prpTDP-43^A315T^ mice upper motor neurons (UMN) showed mitochondrial inner membrane (IMM) integrity loss and endoplasmic reticulum broken cisternae. Interestingly, treatment with the neuroprotective compound NU-9 prevented these ultrastructural abnormalities and ameliorated UMN loss (Genc et al., 2021). A follow up study by the same group used SBT-272, a compound that stabilizes the IMM, to treat TDP-43^A315T^ mice. SBT-272 treatment improved mitochondrial structural integrity and restored mitochondrial motility and function, leading to increased general health of corticospinal motor neurons (Gautam et al., 2023).

Upon inspection of TEM images at low magnification, we found disorganization of axonal microtubules which may contribute to the reduction of synapse number in the transgenic mice. The disorganization of microtubules in *C. elegans* leads to retraction of axons concluding in a reduced number of synaptic boutons (Baran et al., 2010). We also discovered perivascular and peri-astrocytic edema, which suggests alteration in homeostasis of fluids. Neurovascular mechanisms and the disruption of the blood-brain barrier may precede, accelerate, or contribute to chronic disease processes in neurodegenerative disorders of the adult and aging nervous system (Zlokovic, 2008).

Collectively, our findings demonstrate that early TDP-43 mislocalization disrupts synaptic integrity, mitochondrial abundance and cytoskeletal organization in axons. These alterations are consistent with phenotypes observed in ALS and FTD patients. This model allowed us to analyze early changes prior to overt neurodegeneration and provides a platform to study early pathophysiological events key for designing and testing therapeutic strategies (Dejanovic et al., 2024).

## Acknowledgements

We thank members of LANAIS (CONICET) for assisting with TEM mouse tissue processing, Jorge Goldstein for helpful discussion of TEM qualitative assessment, and Veronica Risso and Lucia Garbini for assistance with animal husbandry. We also thank researchers responsible for the public access webtool “TDP-MAP”.

## Authors’ contributions

Conceptualization: FV, LC, LMI; Data curation: FV, ML, JC, LMI; Formal analysis: FV, ML, FLA; Funding acquisition: LMI; Investigation; FV, ML, FLA, JC, LC, LMI; Methodology: FV, JC, LC, LMI; Project administration: LMI; Supervision: LC, LMI; Visualization: FV, ML, JC, LMI; Writing – original draft: FV, JC, LC, LMI; Writing – review and editing: FV, ML, FLA, JC, LC, LMI.

